# An integrated Rheo-NMR platform for real-time, site-specific kinetic analysis of amyloid formation

**DOI:** 10.64898/2026.10.19.760037

**Authors:** Masatomo So, Kei Sato, Kota Yoshioka, Yosuke Shimada, Daichi Morimoto, Hideki Nakamura, Ayako Furukawa, Kenji Sugase

## Abstract

Amyloid fibrils are protein aggregates associated with neurodegenerative diseases. Their formation proceeds through protein– and condition-dependent microscopic pathways that can generate structurally and biologically distinct aggregates, yet these pathways are inferred from ensemble-averaged measurements. Amyloid formation is often slow under quiescent conditions, prompting agitation. However, conventional agitation precludes simultaneous high-resolution solution nuclear magnetic resonance (NMR) spectroscopy, limiting site-specific kinetic analysis of microscopic aggregation pathways. Here we present an integrated Rheo-NMR platform combining controlled shear flow, time-resolved NMR spectroscopy and mechanistic kinetic modeling to quantify site-specific amyloid-formation kinetics in a single, continuously monitored reaction. Applied to α-synuclein under dilute and polyethylene glycol (PEG)-induced liquid-liquid phase separation (LLPS) conditions, the method revealed environment-dependent redistribution of kinetic signatures associated with primary nucleation-elongation and secondary nucleation-elongation pathways across the sequence. The platform distinguishes changes in overall aggregation rates from differences in residue-specific kinetic profiles, providing a quantitative framework for dissecting environment-dependent amyloid formation.

---

The aggregation of proteins into amyloid fibrils is associated with numerous neurodegenerative diseases, including Parkinson’s and Alzheimer’s diseases^1–3^. Structural studies using cryo-electron microscopy (cryo-EM) and solid-state nuclear magnetic resonance (ssNMR) spectroscopy have revealed extensive structural polymorphism among amyloid fibrils. This diversity is observed among fibrils formed under different solution conditions or from disease-associated variants, as well as among fibrils amplified from patient-derived seeds or isolated directly from patient tissues^4–8^. The resulting polymorphs can differ in toxicity, seeding activity and capacity for propagation, potentially contributing to selective cellular vulnerability and variation in disease progression and clinical phenotype. However, these structural methods primarily characterize fibrillar states and provide limited information about the molecular events that precede them.

Understanding how fibrils form and how structural diversity arises therefore requires kinetic analysis of the underlying assembly process. Amyloid formation proceeds through multiple microscopic processes, including primary nucleation, fibril elongation and aggregate-dependent proliferation, whose relative contributions vary among proteins and with the solution environment. Thioflavin T (ThT) fluorescence assays, often combined with mechanistic kinetic modeling, are widely used to infer the relative contributions of these processes from ensemble-averaged aggregation curves^9,10^. However, such analyses cannot determine how individual molecular sites participate in each process. Mutagenesis can identify residues that influence aggregation, but it requires comparisons between separate reactions, and the mutations themselves may alter the conformational ensemble and aggregation mechanism. Defining the molecular events underlying each kinetic process therefore requires site-specific information acquired over time during fibril formation. Solution NMR spectroscopy provides site-specific information by monitoring signals from individual molecular sites^11^. However, in practice, amyloid formation often proceeds too slowly in quiescent NMR samples to be followed within practical measurement times^12,13^. Physical agitation, including shaking and ultrasonication, is widely used to accelerate amyloid formation *in vitro*^14–17^. Because these procedures cannot be applied during conventional high-resolution NMR acquisition, agitation and spectral acquisition need to be performed in separate steps, preventing continuous site-specific observation of aggregation under a defined mechanical environment. Consequently, quantifying site-specific amyloid-formation kinetics within a single continuously monitored aggregation reaction remains difficult.

To address this challenge, we previously developed Rheo-NMR based on a Couette geometry^18^. The apparatus generates controlled shear flow directly within the NMR sample tube, providing mechanical stimulation that promotes amyloid formation while maintaining the magnetic-field homogeneity required for high-resolution spectroscopy^19^. Rheo-NMR therefore enables consecutive acquisition of high-resolution NMR spectra and continuous monitoring of individual NMR signals throughout a single mechanically promoted aggregation reaction. However, the resulting signal trajectories alone cannot quantitatively distinguish the contributions of different microscopic processes. Mechanistic kinetic modeling of these trajectories is therefore needed to quantify site-specific kinetics associated with different amyloid-formation pathways and compare their distributions across solution environments.

Here, we integrated Rheo-NMR with mechanistic kinetic modeling to quantify site-specific amyloid-formation kinetics from time-resolved NMR signals acquired under controlled shear flow. We applied this approach to α-synuclein (αSyn), a disease-associated amyloidogenic protein linked to synucleinopathies, including Parkinson’s disease, multiple system atrophy, and dementia with Lewy bodies^1,20–22^. The aggregation kinetics and fibril structures of αSyn are strongly influenced by the solution environment, and liquid–liquid phase separation (LLPS) can promote its amyloid formation and produce structurally distinct fibrils^23–25^. We compared αSyn amyloid formation under dilute aqueous conditions and polyethylene glycol (PEG)-induced LLPS conditions to examine how the solution environment alters the pre-amyloid conformational ensemble and site-specific fibril-formation kinetics. PEG-induced LLPS altered both the overall rate and the site-specific kinetics associated with the primary and secondary pathways, with changes in the primary pathway linked to the droplet-state ensemble. These results establish Rheo-NMR as a quantitative framework for dissecting how solution environments reshape amyloid-formation mechanisms.

## Results

### Integrated Rheo-NMR workflow for kinetic analysis of amyloid formation

To extract site-specific kinetic parameters during amyloid formation, we established an integrated Rheo-NMR workflow. The workflow begins with preparation of an isotope-labeled sample suited to the molecular sites of interest, followed by selection of a suitable NMR experiment (Fig. 1a). The Rheo-NMR setup generates shear flow in the narrow gap between a stationary inner rod and the rotating NMR tube to promote amyloid formation, which often proceeds slowly under quiescent conditions (Fig. 1b). The selected NMR experiment, such as a two-dimensional heteronuclear single-quantum coherence (HSQC) experiment, is then repeated throughout aggregation to monitor molecular changes in real time (Fig. 1c). Time-dependent chemical shift changes or intensity decays are extracted for individual NMR signals associated with specific molecular sites (Fig. 1d). The same workflow can be applied under different solution conditions to compare their effects on amyloid formation kinetics.

**Fig. 1.**
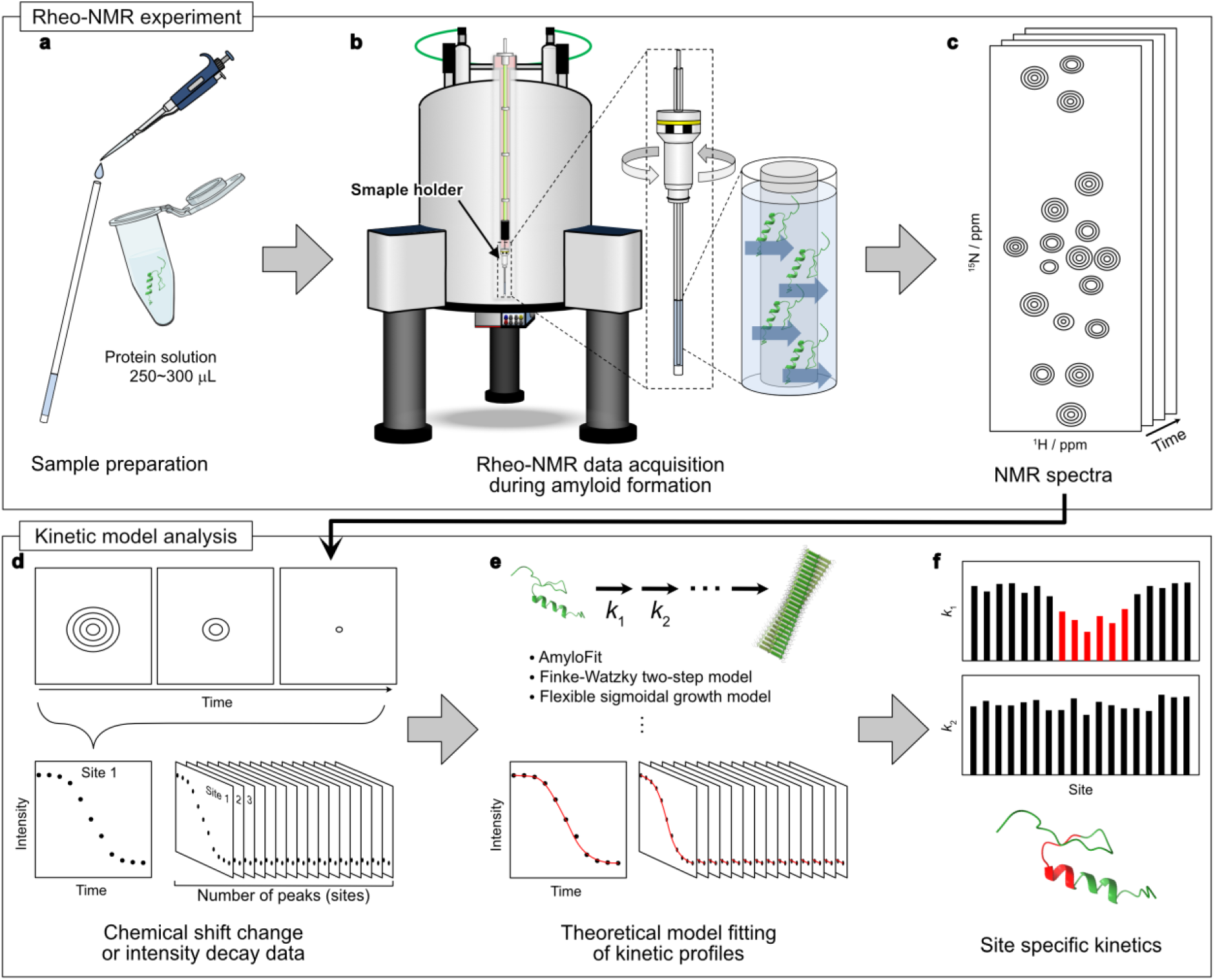
| Integrated Rheo-NMR workflow for site-specific kinetic analysis of amyloid formation. **a–c**, Experimental setup and real-time data acquisition. **a**, Protein solution (250– 300 μL) is placed in an NMR tube. **b**, Controlled shear flow is generated in the sample within the narrow gap between a stationary inner rod and the rotating NMR tube. **c**, NMR spectra are acquired consecutively during amyloid formation to generate a time series, illustrated here using two-dimensional spectra. **d–f**, Extraction of signal-specific kinetic parameters. **d**, Time-course profiles of chemical shift changes or signal-intensity decays are obtained for each resolved NMR signal. **e**, The profiles are globally fitted to an appropriate kinetic model to extract signal-specific kinetic parameters. **f**, The extracted kinetic parameters are plotted as a function of molecular site and mapped onto the protein structure to visualize site-specific differences in amyloid-formation kinetics.

A variety of kinetic models have been developed and applied to ensemble-averaged time-course data obtained from conventional assays such as ThT fluorescence. These include the Finke–Watzky two-step model^26^, a flexible sigmoidal growth model^27^, and mechanistic nucleation-and-growth models implemented in AmyloFit^28^. Applying an appropriate model to Rheo-NMR data allows kinetic parameters to be determined from the time-course profile of each NMR signal. Global fitting of multiple signal-specific profiles, with some parameters shared across all signals and others allowed to vary among signals, yields both global and signal-specific kinetic parameters (Fig. 1e). Comparing these parameters across molecular sites and mapping them onto the protein structure reveals regions with distinct kinetic behavior during amyloid formation, providing molecular-level insight into the aggregation mechanism (Fig. 1f).

### Application of the integrated Rheo-NMR workflow to αSyn amyloid formation

We next applied the integrated Rheo-NMR workflow to αSyn under dilute conditions to investigate residue-specific kinetics during amyloid fibril formation. Under quiescent conditions, the ^1^H-^15^N HSQC spectra of αSyn remained unchanged for 3 days, indicating that no detectable aggregation occurred during this period (Extended Data Fig. 1). In contrast, rotation at 20 Hz in the Rheo-NMR setup, corresponding to an estimated shear rate of approximately 400 s^-1^, induced amyloid fibril formation within 30 h. Amyloid fibril formation was confirmed by a marked increase in ThT fluorescence, the development of visible turbidity, and fibrillar structures observed by TEM (Fig. 2a–c). In the corresponding time-resolved HSQC spectra, peak positions remained essentially unchanged, whereas peak intensities showed little change during an initial lag phase and then decreased sigmoidally (Fig. 2d,e and Extended Data Figs. 2–6). The resulting intensity-decay profiles, which reflect the loss of NMR-visible αSyn during amyloid fibril formation, varied among residues, enabling residue-specific kinetic analysis.

**Fig. 2.**
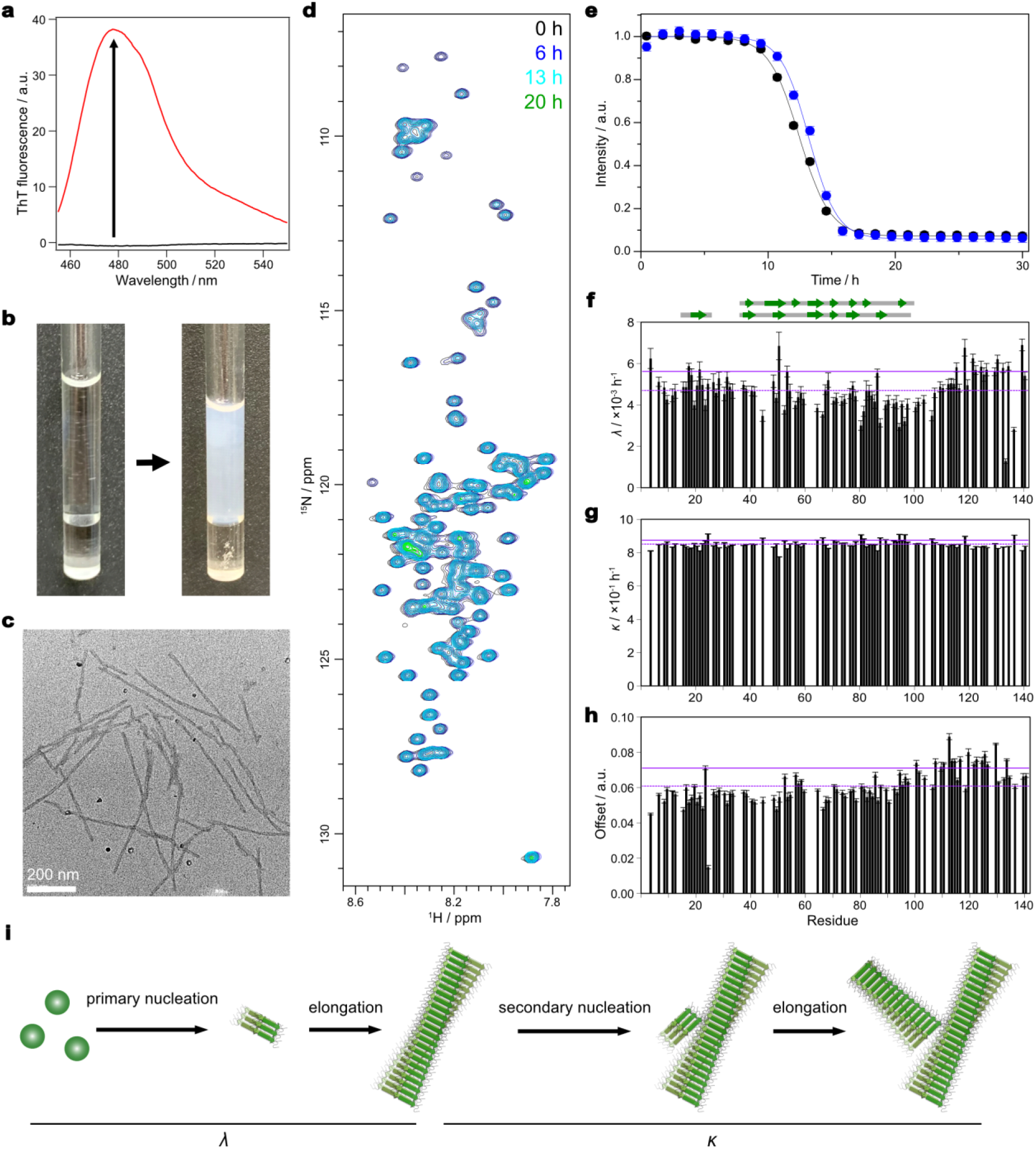
| Rheo-NMR analysis of αSyn amyloid fibril formation under dilute conditions. **a**, Thioflavin T (ThT) fluorescence spectra before (black) and after (red) 30 h of rotation. **b**, Images of the sample before (left) and after (right) 30 h of rotation. **c**, TEM image of the sample after 30 h of rotation. Scale bar, 200 nm. **d**, ^1^H-^15^N HSQC spectra of αSyn acquired at 0, 6, 13, and 20 h during 20-Hz rotation at 30 °C. **e**, Time courses of HSQC peak intensities for Asp121 (black) and Tyr136 (blue), normalized to the values before rotation. Solid lines indicate the best-fit curves. Error bars represent fitting errors. Profiles for all assigned residues are shown in Extended Data Figs. 2–6. **f–h**, Residue-specific fitted values of the composite kinetic parameters *λ* (**f**) and *κ* (**g**), and the residual intensity *I*_offset_ (**h**). Purple dotted and solid lines represent the mean and the mean plus one standard deviation, respectively. Error bars represent standard deviations obtained from the fitting. β-strand and turn regions in representative αSyn fibril structures from the two major classes described by Milchberg et al.^33^ are shown above panel **f** (PDB entries 6XYQ^8^ and 6RT0^34^). **i**, Schematic representation of the modified closed-form secondary-nucleation model.

To quantify the residue-specific kinetics, we fitted the intensity-decay profiles with a modified closed-form secondary-nucleation model^28^ (Fig. 2i). The use of this model is supported by recent studies identifying secondary nucleation as the dominant mechanism of αSyn fibril amplification under specific physicochemical conditions^29^. The analysis yielded two composite kinetic parameters, *λ* and *κ* (Fig. 2f,g). *λ* combines primary nucleation with elongation, whereas *κ* combines monomer-dependent secondary nucleation with elongation. The fitted *λ* and *κ* values were on the order of 10^-3^ and 10^-1^ h^-1^, respectively, consistent with the slow generation of initial fibrils through the primary pathway followed by rapid amplification through the secondary pathway. The *λ* values were lower in the NAC region than in the N– and C-terminal regions, whereas the *κ* values were nearly uniform across the sequence. The lower *λ* values in the NAC region, which forms a major part of the structural core of mature αSyn fibrils, suggest that αSyn amyloid fibril formation proceeds through a stepwise primary pathway initiated by intermolecular association of the flexible N– and C-terminal regions, followed by structural rearrangement of the initial assemblies into a NAC-region-centered amyloid core and subsequent fibril elongation. By contrast, the uniform *κ* values suggest that secondary-nucleation-driven amplification involves the N-terminal, NAC, and C-terminal regions with similar kinetics, rather than progressing stepwise from one region to another.

The fitted *I*_offset_ values were nonzero (Fig. 2h), likely reflecting contributions from monomeric αSyn and/or mobile regions in oligomeric or fibrillar species. Importantly, *I*_offset_ was higher in the C-terminal region than in the N-terminal and NAC regions. This region-specific persistence is consistent with the C-terminal region remaining relatively mobile and NMR-visible in oligomeric or fibrillar species^30–32^.

### Amyloid formation under PEG-induced LLPS conditions

Previous studies have shown that PEG induces the formation of αSyn liquid droplets, which subsequently convert into amyloid fibrils^23,35^. Cryo-EM analysis has also resolved the structure of αSyn fibrils formed from liquid droplets^25^. To examine molecular events associated with environment-dependent fibril polymorphism, we first characterized αSyn under PEG-induced LLPS conditions without flow. Before the NMR measurements, fluorescence microscopy confirmed the formation of liquid droplets in the presence of PEG, whereas no droplets were observed in its absence (Fig. 3a), consistent with previous reports^23,36^. We then compared the ^1^H-^15^N HSQC spectra recorded in the presence and absence of PEG under quiescent conditions (Fig. 3b). Residues showing large chemical shift perturbations (CSPs) were distributed across the sequence but were particularly frequent within the fibril-core region (residues 38–97) and were also found in the N– and C-terminal regions (Fig. 3c). The observed CSPs indicate changes in αSyn conformation, intermolecular interactions, or both. Mapping residues with large CSPs onto the αSyn fibril structure formed from liquid droplets^25^ revealed spatial clustering (Fig. 3d), suggesting a possible relationship between the pre-amyloid αSyn ensemble and the reported fibril structure. After incubation at 25 °C under quiescent conditions for 5 days, the HSQC spectrum showed only minor changes (Extended Data Fig. 7), suggesting that αSyn remained stable for at least 5 days even in the presence of PEG.

**Fig. 3.**
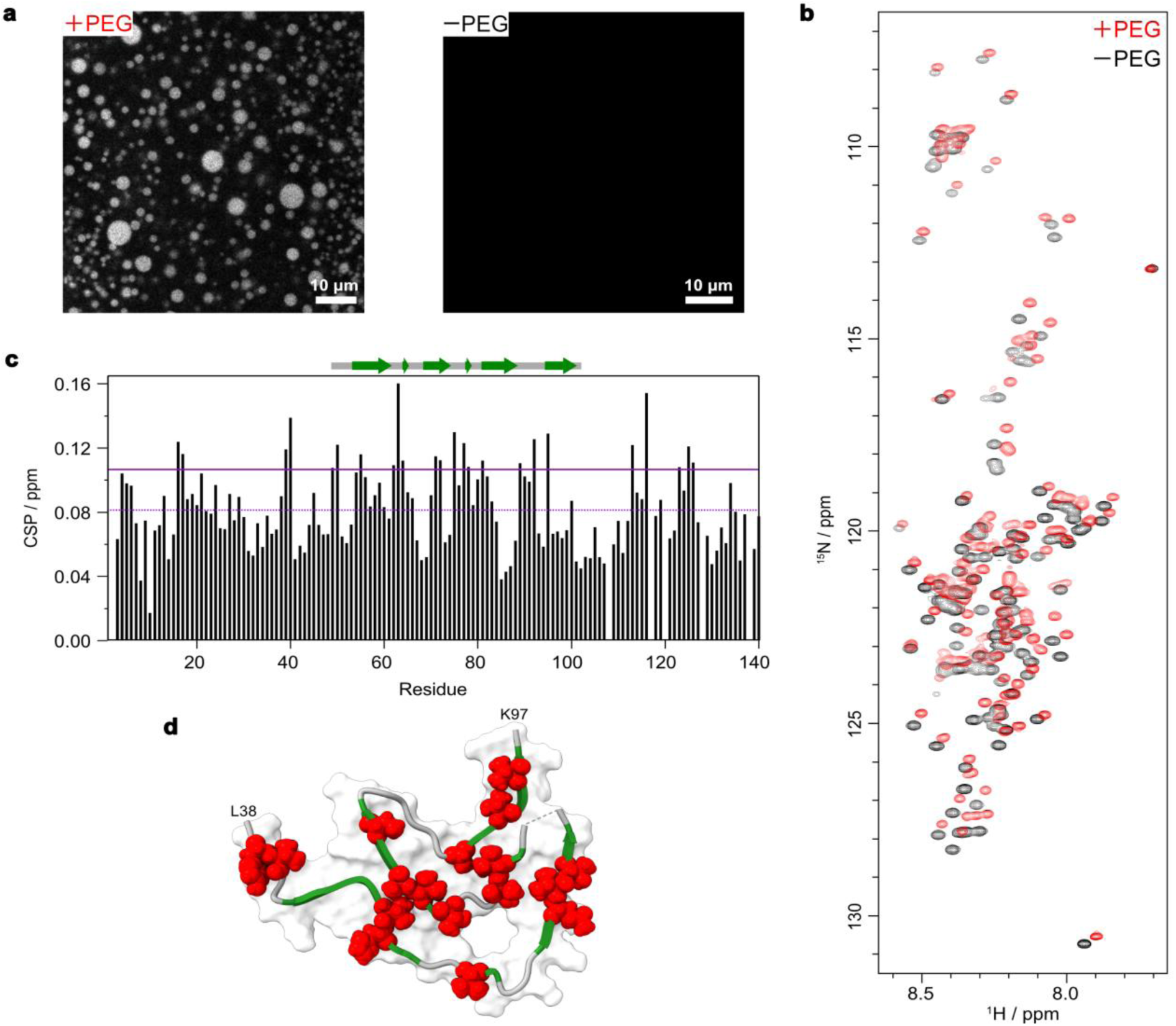
| NMR analysis of αSyn in the presence of PEG under quiescent conditions. **a**, Fluorescence microscopy images of αSyn in the presence (left) and absence (right) of PEG, acquired before the NMR measurements. Scale bars, 10 μm. **b**, ^1^H-^15^N HSQC spectra of αSyn in the presence (red) or absence (black) of PEG under quiescent conditions at 25 °C. **c**, Residue-specific chemical shift perturbations (CSPs) induced by PEG. Purple dotted and solid lines indicate the mean and the mean plus one standard deviation, respectively. β-strand and turn regions observed in the αSyn fibril structure formed from liquid droplets (PDB entry 8RI9)^25^ are shown in green and gray, respectively, above the plot. **d**, Residues with CSPs exceeding the mean plus one standard deviation were mapped onto the αSyn fibril structure formed from liquid droplets (PDB entry 8RI9)^25^ using UCSF ChimeraX^37^ and are shown as red spheres.

We next subjected this αSyn sample to 20-Hz rotation during Rheo-NMR measurements under the same PEG-induced LLPS conditions. αSyn amyloid fibrils formed within 13 h, faster than under dilute conditions, as confirmed by a marked increase in ThT fluorescence, the appearance of visible white aggregates, and fibrillar structures observed by TEM (Fig. 4a–c). In the corresponding time-resolved HSQC spectra, peak positions remained unchanged, whereas peak intensities decreased sigmoidally (Fig. 4d,e and Extended Data Figs. 8–10). The resulting intensity-decay profiles again varied among residues.

**Fig. 4.**
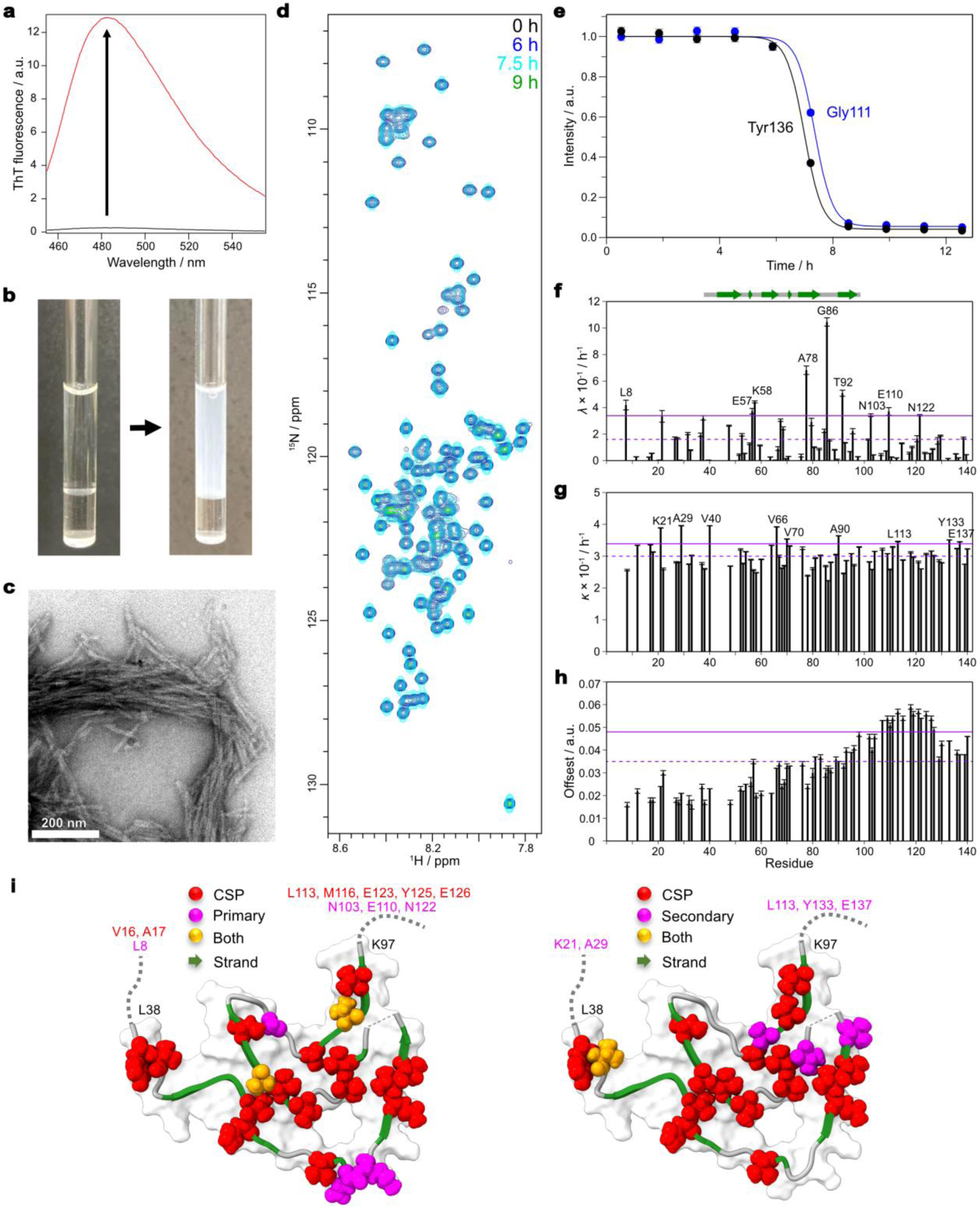
| Rheo-NMR analysis of αSyn amyloid fibril formation under PEG-induced LLPS conditions. **a**, Thioflavin T (ThT) fluorescence spectra before (black) and after (red) 13 h of rotation. **b**, Images of the sample before (left) and after (right) 13 h of rotation. **c**, TEM image of the sample after 13 h of rotation. Scale bar, 200 nm. **d**, ^1^H-^15^N HSQC spectra of αSyn under PEG-induced LLPS conditions at 0, 6, 7.5, and 9 h during 20-Hz rotation at 25 °C. **e**, Time courses of HSQC peak intensities for Tyr136 (black) and Gly111 (blue), normalized to the values before rotation. Solid lines indicate the best-fit curves. Error bars represent fitting errors but are smaller than the symbols. Profiles for all assigned residues are shown in Extended Data Figs. 8–10. **f–h**, Residue-specific fitted values of the composite kinetic parameters *λ* (**f**) and *κ* (**g**), and the residual intensity *I*_offset_ (**h**). Purple dotted and solid lines represent the mean and the mean plus one standard deviation, respectively. Error bars represent standard deviations obtained from the fitting. β-strand and turn regions in the αSyn fibril structure formed from liquid droplets (PDB entry 8RI9)^25^ are shown as green arrows and gray bars, respectively, above panel **f**. **i**, Residues with CSPs exceeding the mean plus one standard deviation (red), *λ* values exceeding the mean plus one standard deviation (purple), or both (orange) are mapped onto the fibril structure formed from liquid droplets (PDB entry 8RI9)^25^ in the left panel. Residues with CSPs exceeding the mean plus one standard deviation (red), *κ* values exceeding the mean plus one standard deviation (purple), or both (orange) are shown on the same structure in the right panel. β-strands are represented by green arrows.

To quantify the residue-specific kinetics, we again fitted the intensity-decay profiles with the modified closed-form secondary-nucleation model, yielding residue-specific *λ* and *κ* values (Fig. 4f,g). The residue-specific *λ* profile differed between conditions. Under PEG-induced LLPS conditions, high *λ* values were prominent within and around the fibril-core region (Fig. 4f), particularly at sites exhibiting large CSPs under quiescent conditions (Fig. 4i, left), whereas under dilute conditions they were observed mainly in the N– and C-terminal regions (Fig. 2f). Residues exhibiting both high *λ* values and large CSPs included E57 and K58 within the β-arc lineage region (G51–G67), as well as A78, G86, and T92 within the Greek-key lineage region (A69–A95) (Fig. 4i, left). This overlap suggests a residue-level association between the pre-amyloid αSyn ensemble and the primary pathway represented by *λ*, potentially biasing subsequent fibril assembly. By comparison, high *κ* values under PEG-induced LLPS conditions were distributed broadly across the sequence, with local maxima in the flexible terminal regions and at several loop or turn sites within the fibril core (Fig. 4i, right). Such heterogeneity was not observed under dilute conditions, where the *κ* profile was nearly uniform (Fig. 2g).

By contrast, the fitted *I*_offset_ values under PEG-induced LLPS conditions were again nonzero, with higher values in the C-terminal region (Fig. 4h). This common pattern is consistent with the C-terminal region remaining relatively mobile and NMR-visible in oligomeric or fibrillar species under both conditions. Together, these results indicate that the solution environment modulates not only the rate of αSyn amyloid fibril formation but also the residue-specific kinetic profiles of the primary and secondary pathways, with potential implications for fibril polymorphism.

## Discussion

Here, we established an integrated Rheo-NMR platform that combines controlled shear flow with time-resolved NMR spectroscopy and kinetic modeling to quantify residue-specific amyloid formation kinetics in real time (Fig. 1). Application of this platform to αSyn revealed distinct residue-specific kinetic signatures associated with the primary and secondary pathways under dilute and PEG-induced LLPS conditions (Figs. 2 and 4). These differences demonstrate that the solution environment reshapes the residue-specific kinetics of αSyn amyloid formation and may thereby influence fibril polymorphism. Rheo-NMR thus provides mechanistic information that is not accessible from conventional ensemble-averaged kinetic measurements alone.

The Rheo-NMR setup uses a Couette geometry to generate controlled shear flow, promoting amyloid formation while maintaining the magnetic-field homogeneity required for the consecutive acquisition of high-resolution NMR spectra^18,19^. Reliable Rheo-NMR measurements require a stable and well-defined shear field, for which precise coaxial alignment of the inner glass rod and outer NMR tube is essential. In this regard, the recently developed motor-driven Rheo-NMR system mechanically integrates the inner and outer components within a unified sample-cell assembly that precisely fixes their coaxial and vertical alignment^38^, thereby improving mechanical stability and the consistency of the applied shear conditions.

The kinetic model includes primary nucleation, elongation and monomer-dependent secondary nucleation^28^, consistent with previous studies showing that αSyn aggregates proliferate through secondary nucleation and other aggregate-dependent processes^29,39–41^. Combining Rheo-NMR with this kinetic model extends established kinetic analyses of ensemble-averaged aggregation data to residue-specific profiles obtained within a single reaction. Unlike mutation-based ThT studies, which infer residue contributions from separate measurements of variant proteins^42–46^, Rheo-NMR provides kinetic information for multiple residues across wild-type αSyn within a single reaction, without introducing mutation-dependent changes in the conformational ensemble or aggregation environment.

Under dilute conditions, the N– and C-terminal regions showed higher *λ* values than the NAC region (Fig. 2f). This may appear counterintuitive because the NAC region forms a major part of the rigid core of mature fibrils. However, the observed pattern is consistent with studies showing that short N-terminal motifs regulate the early stages of amyloid formation and oligomer-to-fibril conversion^42,43,46^ and that the N terminus mediates recruitment of soluble αSyn to fibrils^31,32^. The C-terminal tail is also critical for oligomerization while suppressing aggregate replication^45^, and His50 modulates αSyn aggregation at physiological pH^44^. Together, these findings suggest a stepwise primary pathway initiated by intermolecular association of the flexible terminal regions, followed by structural rearrangement into a NAC-region-centered amyloid core and fibril elongation. In contrast to the regional variation in *λ*, *κ* was nearly uniform across the sequence (Fig. 2g), suggesting broader sequence involvement once aggregate-dependent proliferation begins.

Given the structural diversity of αSyn fibrils *in vitro* and in human disease^8,34,47–49^, we examined whether changes in the solution environment alter the pre-amyloid conformational ensemble and residue-specific fibril-formation kinetics in ways that may influence fibril structure. Under quiescent conditions with PEG, residues showing large CSPs clustered spatially when mapped onto the reported fibril structure formed from liquid droplets^25^ (Fig. 3d), suggesting a relationship between the pre-amyloid αSyn ensemble and this fibril fold. PEG-induced LLPS did not simply accelerate fibril formation but also altered the residue-specific *λ* and *κ* profiles, with high *λ* sites overlapping residues showing large CSPs (Figs. 2f,g and 4f,g,i). This overlap suggests a residue-level link between the pre-amyloid ensemble and subsequent fibril-formation kinetics. The faster fibril formation under PEG-induced LLPS conditions is also consistent with previous reports of nucleation and aggregate-dependent proliferation within αSyn condensates^23–25,36,40^. Previous studies under dilute conditions have identified long-range contacts involving the terminal and central hydrophobic regions and linked N-terminal exposure to aggregation propensity^50–53^, whereas the CSP pattern under PEG-induced LLPS conditions is consistent with the formation of local hydrophobic clusters involving residues in the fibril-core region. Together, these findings support a model in which environment-dependent changes in the pre-amyloid ensemble may bias early fibril assembly and thereby influence fibril polymorphism, while secondary processes promote fibril proliferation (Fig. 5).

**Fig. 5.**
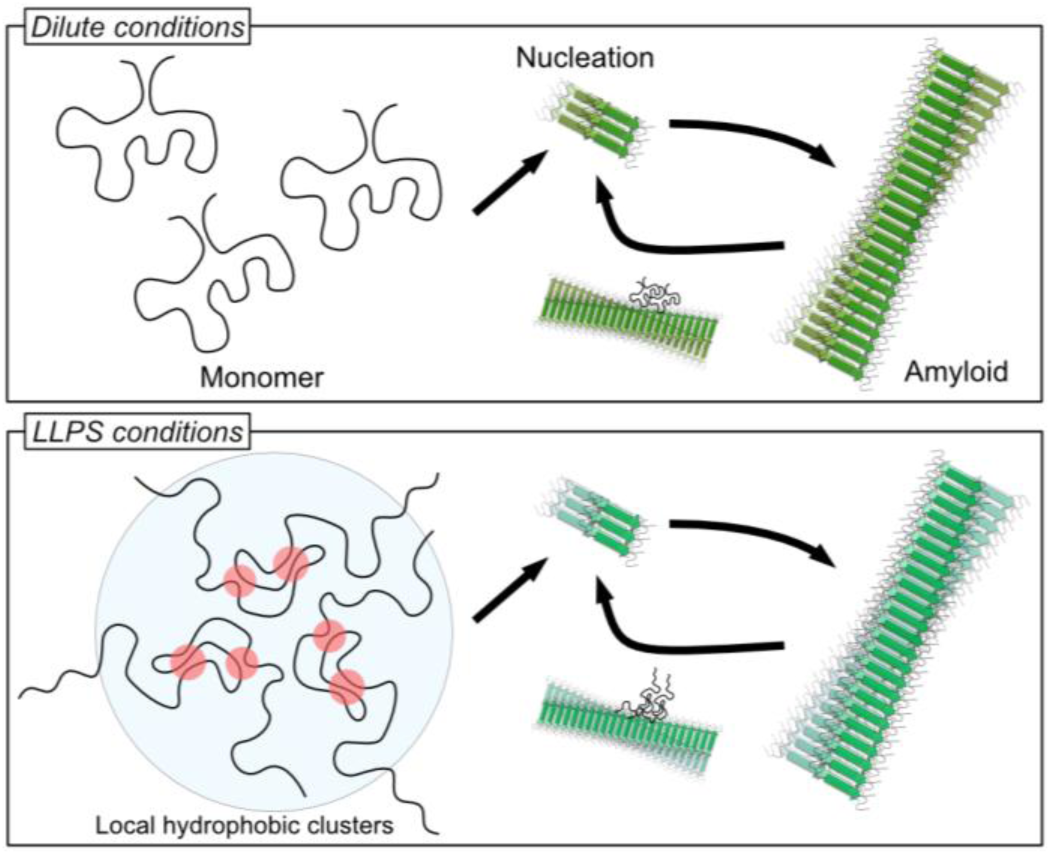
| Working model for environment-dependent αSyn amyloid formation. Schematic model of αSyn amyloid formation under dilute conditions (upper) and PEG-induced LLPS conditions (lower).

Although applying a kinetic model developed for ensemble-averaged aggregation data to residue-specific NMR signal changes requires careful interpretation, the approach resolves local kinetic differences across the αSyn sequence. Global analysis across a series of initial protein concentrations could further constrain the reaction orders and deconvolute the rate-constant combinations underlying *λ* and *κ*^28^. Such measurements are more challenging in Rheo-NMR than in ThT assays because reducing protein concentration weakens NMR signals, whereas the detection sensitivity of ThT assays can, to some extent, be adjusted through the dye concentration. More generally, Rheo-NMR requires sufficient sensitivity and spectral resolution, as well as acquisition times short enough to follow aggregation. In the present study, αSyn required 30 °C to form fibrils without PEG, whereas fibrillation with PEG at 30 °C was too rapid for the temporal resolution of the NMR measurements and was therefore monitored at 25 °C. This limitation could be addressed by higher-sensitivity NMR measurements, which would permit shorter acquisition times and temperature-matched comparisons.

More broadly, the isotope-labeling scheme, NMR experiment, solution conditions and kinetic model can be adapted to diverse amyloidogenic proteins. Systematic variation of pH, ionic strength, molecular crowding, membrane environments and small-molecule additives could determine how specific solution factors redistribute pathway-associated kinetics and potentially influence fibril polymorphism. In a complementary study, Rheo-NMR revealed preferential retention of oxidized epigallocatechin gallate–αSyn interactions during shear-promoted assembly^54^. The same framework could be extended to other phase-separating systems to determine how different environmental triggers reshape the transition from soluble states to fibrils. Applying this approach across proteins and environments may help develop mechanistic models of amyloid formation, clarify how pathological aggregation begins and identify molecular interactions and kinetic processes that could serve as targets for developing and evaluating fibril-formation inhibitors.

In conclusion, we established a platform that integrates Rheo-NMR with mechanistic kinetic modeling to enable the real-time, residue-specific analysis of amyloid formation. This approach resolves the trade-off between the mechanical acceleration required to promote aggregation and the magnetic-field homogeneity required for high-resolution NMR. Application to αSyn revealed both the environment-dependent redistribution of kinetic signatures associated with the primary and secondary pathways across the sequence and a residue-level correspondence between pre-amyloid NMR perturbations and subsequent fibril-formation kinetics. The integrated platform thus provides a general framework for investigating how protein-specific properties and the solution environment shape amyloid formation and fibril polymorphism.

## Data availability

All data supporting the findings of this study are available within the article, its Extended Data and the accompanying Source Data file. The resonance assignments and fibril structure used in this study are available under BMRB accession 6968 and PDB accession 8RI9, respectively. Additional data are available from the corresponding author upon reasonable request.

## Code availability

The customized version of GLOVE used for kinetic fitting in this study is available from the corresponding author upon reasonable request.

## Acknowledgments

This work was supported by JSPS KAKENHI (Grant Numbers 17K19196, 18H04551, 18KK0075, 21H02442, 22H05088, 22H04842, 25K00268 and 25K02240), the Japan Keirin Association grants (JKA; grant numbers 2021M-224, 2023M-363, 2024M-517, and 2025M-491), the Kyoto University 125th Anniversary Fund “Kusunoki 125”, the Toyota Riken Scholar and the Toyota Riken Scholar Joint Research of the Toyota Physical and Chemical Research Institute, the Mitsubishi Foundation, the Hachiro Honjo Ocha Foundation, and the Takeda Science Foundation. This work was performed in part under the Cooperative Research Program and the Collaborative Research Program of Institute for Protein Research, Osaka University (CR-25-02 and NMRCR-25-05). TEM observations were performed at the North Campus Instrumental Analysis Station, Kyoto University.

## Author contributions

K. Sato, K.Y., Y.S., M.S., and H.N. performed experiments and analyzed the data. M.S. wrote the initial draft of the manuscript. M.S., D.M., H.N., and A.F. contributed to manuscript revision. K. Sugase conceived and designed the study, acquired funding, supervised the project, contributed to data interpretation and discussion, and wrote and finalized the manuscript. All authors reviewed and approved the final manuscript.

## Additional information

### Extended data

Extended Data Figs. 1–10 are available with the online version of this paper.

## Methods

### Expression and purification of αSyn

Recombinant human αSyn bearing an N-terminal GST-SUMO tag was expressed and purified as described previously. Briefly, the fusion protein was expressed in *E. coli* BL21 cells and purified by glutathione-affinity chromatography using a GSTrap 4B column (Cytiva). The GST-SUMO tag was cleaved with SENP2 protease, and the resulting untagged αSyn was purified again by glutathione-affinity chromatography followed by anion-exchange chromatography using a HiTrap Q HP column (Cytiva). The molecular mass of αSyn was confirmed using an autoflex MALDI-TOF mass spectrometer (Bruker).

### NMR measurements under quiescent conditions

Lyophilized [*U*-^15^N]-αSyn was dissolved in 6 M guanidine hydrochloride. The sample was buffer-exchanged into 40 mM MOPS (pH 7.0) using a Zeba Spin desalting column (Thermo Fisher Scientific). The αSyn concentration was adjusted to 500 µM in 40 mM MOPS (pH 7.0) containing 150 mM NaCl and 5% D_2_O, either without or with 10% (w/v) PEG 8000. For measurements under quiescent conditions, ^1^H-^15^N HSQC spectra were acquired at 25 °C. Backbone amide resonance assignments were transferred from BMRB entry 6968 and verified by confirming carbonyl-carbon connectivities using the MOCCA XY-16 isotropic mixing pulse sequence. Residue-specific CSPs induced by PEG were calculated as Δ*δ* = [(Δ*δ*_H_)^2^ + (Δ*δ*_N_/5)^2^]^1/2^, where Δ*δ*_H_ and Δ*δ*_N_ are the chemical shift differences in the ^1^H and ^15^N dimensions, respectively. Sample stability in the presence of PEG was assessed by comparing ^1^H-^15^N HSQC spectra acquired before and after 5 days of quiescent incubation at 25 °C. NMR data were processed using NMRPipe, and the spectra analyzed using CcpNmr AnalysisAssign.

### Rheo-NMR measurements of αSyn amyloid fibril formation

Rheo-NMR measurements were performed using a previously described cryogenic-probe-compatible setup on AVANCE I 600 or AVANCE II 700 NMR spectrometers, each equipped with a *z*-gradient TCI cryogenic probe (Bruker). An NMR tube containing 250 µL of the sample solution was placed in the NMR magnet without a cap. The µRheo-NMR device (AirPro) was then inserted into the NMR tube. The NMR tube was rotated at 20 Hz using the sample-spinning system of the spectrometer to generate shear flow. Two-dimensional ^1^H-^15^N heteronuclear single-quantum coherence (HSQC) spectra were acquired consecutively with a temporal resolution of 1.29 h per spectrum under shear flow at 30 °C without PEG and at 25 °C with 10% (w/v) PEG 8000 to monitor αSyn amyloid fibril formation in real time. Rheo-NMR spectra were processed using TopSpin (Bruker), and peak intensities were analyzed using PINT.

### Kinetic analysis of NMR peak intensities

For each residue, the time-dependent peak intensity, *I*(t), was fitted with a customized version of GLOVE to a modified closed-form integrated rate law for polymer mass accumulation in a model comprising primary nucleation, elongation, and monomer-dependent secondary nucleation, as described previously^28^. The original rate law describes the increase in polymer mass fraction, whereas the decrease in HSQC peak intensity was used here to monitor the loss of NMR-visible αSyn. Assuming that the peak intensity was proportional to the amount of NMR-visible αSyn, the decay of the NMR-visible component was described by the complement of the polymer mass fraction, with an additive offset (*I*_offset_) representing the asymptotic residual peak intensity, as follows:

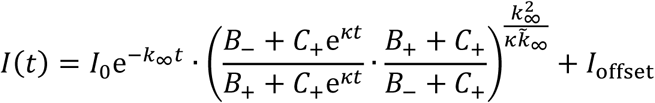

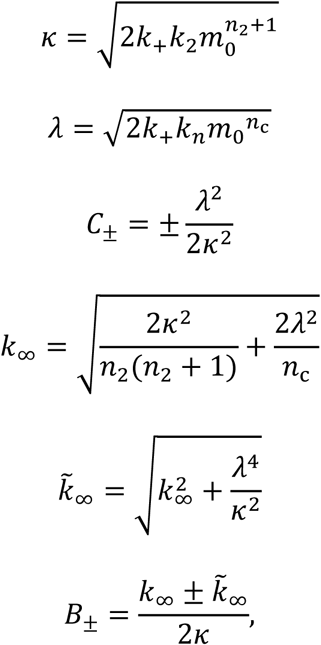

where *I*_0_ is the initial intensity of the decaying NMR-visible component; *k*_+_, *k*_n_, and *k*_2_ are the rate constants for elongation, primary nucleation, and monomer-dependent secondary nucleation, respectively; *n*_c_ and *n*_2_ are the reaction orders for primary nucleation and monomer-dependent secondary nucleation, respectively; and *m*_0_ is the initial monomer concentration. *λ* combines primary nucleation with elongation, whereas *κ* combines monomer-dependent secondary nucleation with elongation. Both *n*_c_ and *n*_2_ were treated as global parameters across all residues, whereas *I*_0_, *I*_offset_, *λ*, and *κ* were optimized independently for each residue.

### Thioflavin T fluorescence assay

A 1-µL aliquot of the sample solution was added to 1 mL of the corresponding buffer containing 25 µM thioflavin T (ThT). ThT fluorescence emission spectra were recorded on a FluoroMax-4 spectrofluorometer (HORIBA) with excitation at 445 nm.

### Transmission electron microscopy

The sample solution was diluted tenfold, and 5 μL of the diluted solution was spotted onto a collodion-coated copper grid (Nisshin EM). After 1 min, excess solution was blotted away with filter paper, and 5 μL of EM Stainer solution (Nisshin EM) was spotted onto the grid. Excess stain was removed in the same manner, and the grid was rinsed with Milli-Q water. Transmission electron microscopy (TEM) images were acquired using a JEM-1400Plus transmission electron microscope (JEOL) operated at an accelerating voltage of 80 kV over a magnification range of 10,000–30,000.

### Fluorescence microscopy

To visualize protein condensation by fluorescence microscope, α-Syn protein was fluorescently labeled with NHS-rhodamine via NHS ester conjugation to primary amines. Samples containing 10% rhodamine-labeled αSyn were placed on Pluronic F-127-treated glass slides and immediately imaged using a DMi8 fluorescence microscope equipped with a 63× oil-immersion objective (Leica Microsystems).

