## Extended Data for "An integrated Rheo-NMR platform for real-time, site-specific kinetic analysis of amyloid formation"

**Table of Contents**

Extended Data Figs. 1–10

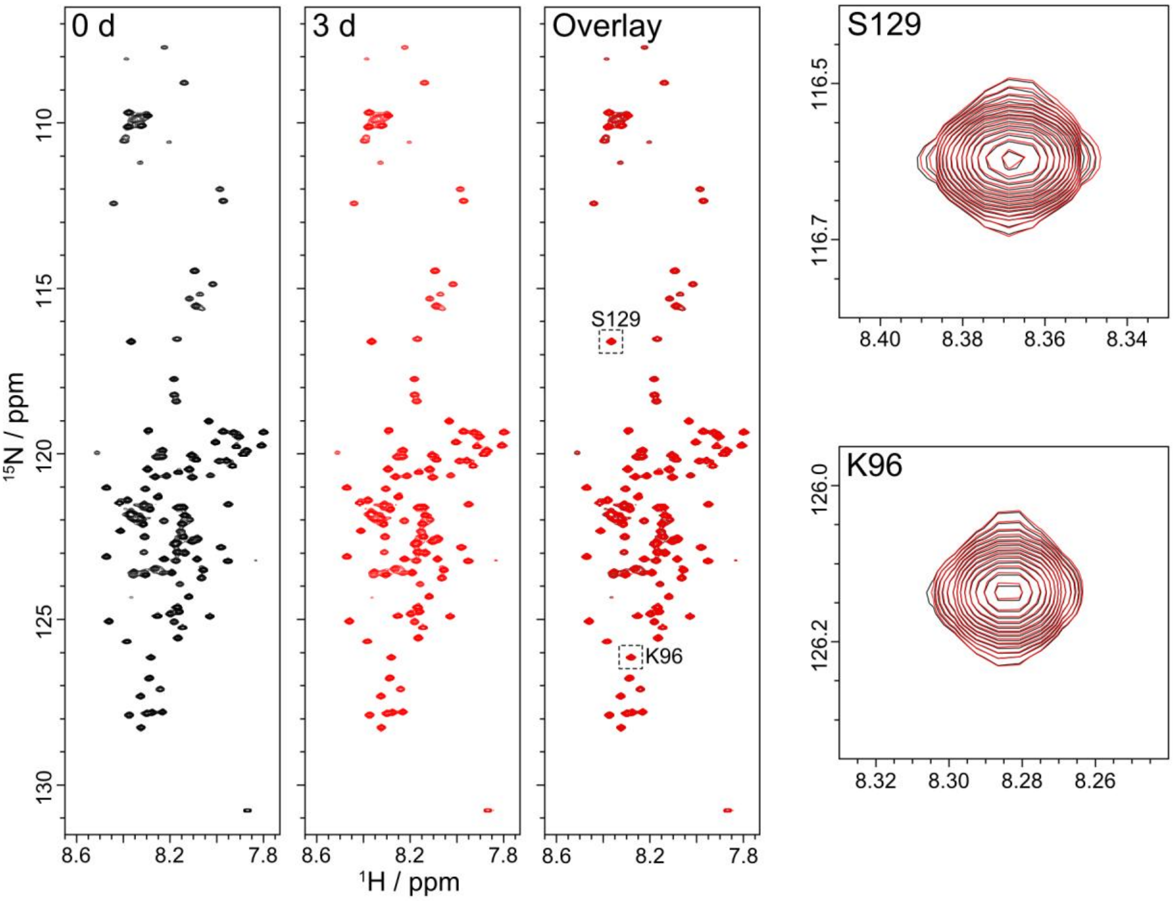


**Extended Data Fig. 1 | NMR spectra under quiescent conditions.** $^{1}$H-$^{15}$N HSQC spectra of αSyn acquired before (black) and after 3 days (red) of incubation at 25 °C. Signals from Lys96 and Ser129 are enlarged to show their complete overlap.

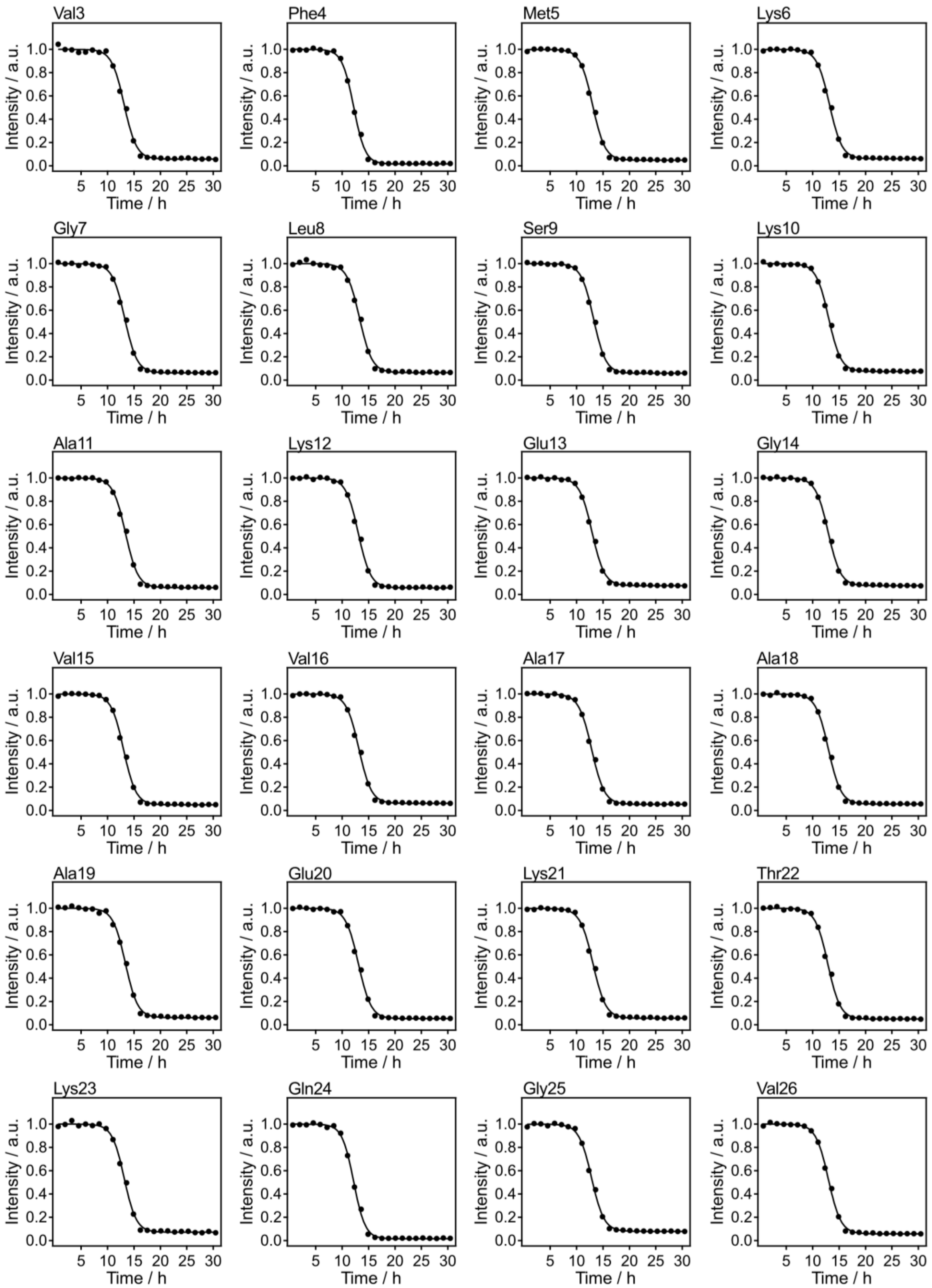


**Extended Data Fig. 2 | Rheo-NMR measurements of αSyn amyloid fibril formation under dilute conditions.** Time courses of $^{1}H$-$^{15}N$ HSQC peak intensities for assigned residues from Val3 to Val26 during 20-Hz rotation at 30 °C without PEG. Intensities were normalized to the values before rotation. Solid lines indicate the best-fit curves obtained using the equation shown in Methods. Error bars represent fitting errors.

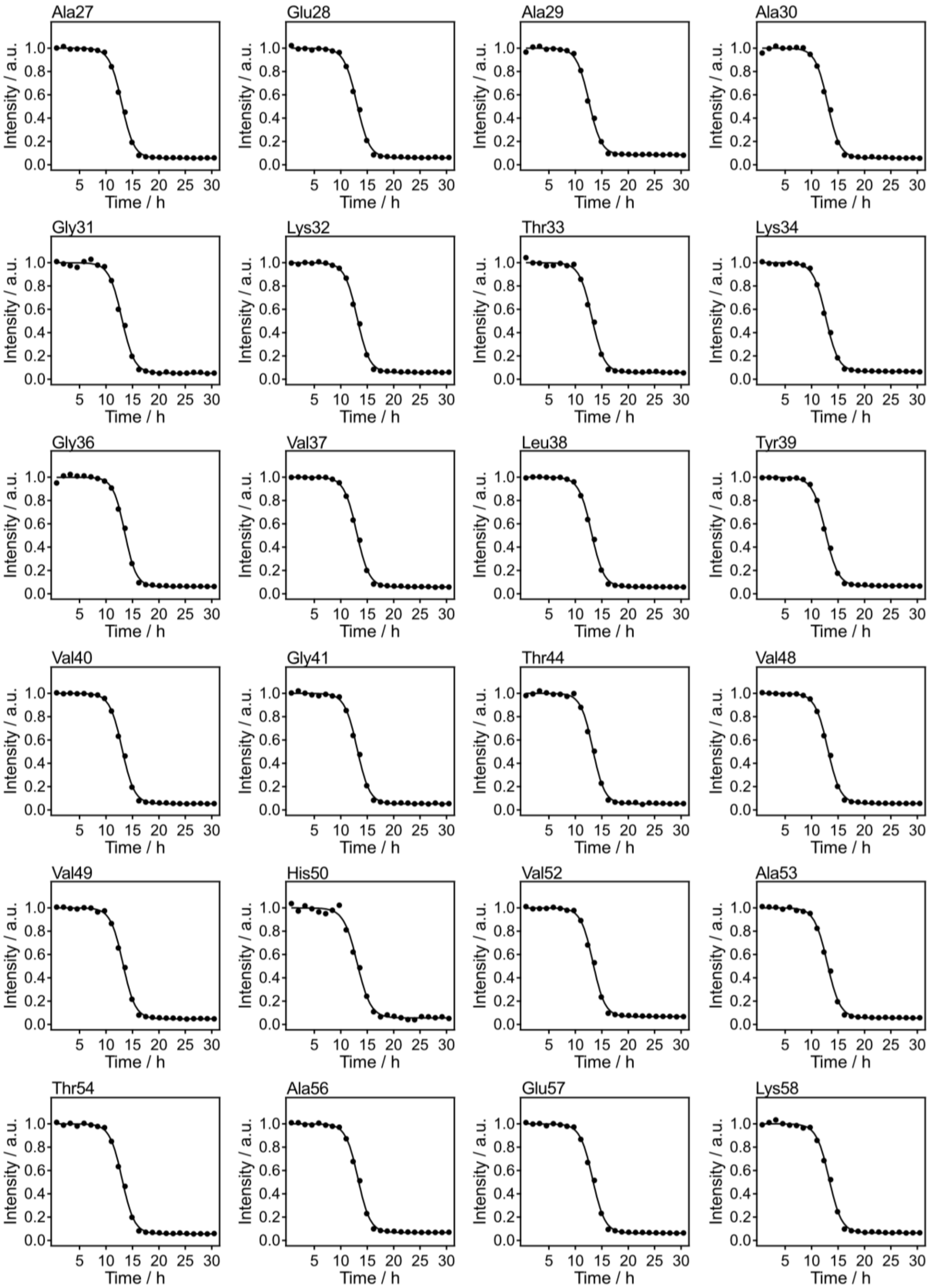


**Extended Data Fig. 3 | Rheo-NMR measurements of αSyn amyloid fibril formation under dilute conditions.** Time courses of $^{1}H$-$^{15}N$ HSQC peak intensities for assigned residues from Ala27 to Lys58 during 20-Hz rotation at 30 °C without PEG. Intensities were normalized to the values before rotation. Solid lines indicate the best-fit curves obtained using the equation shown in Methods. Error bars represent fitting errors.

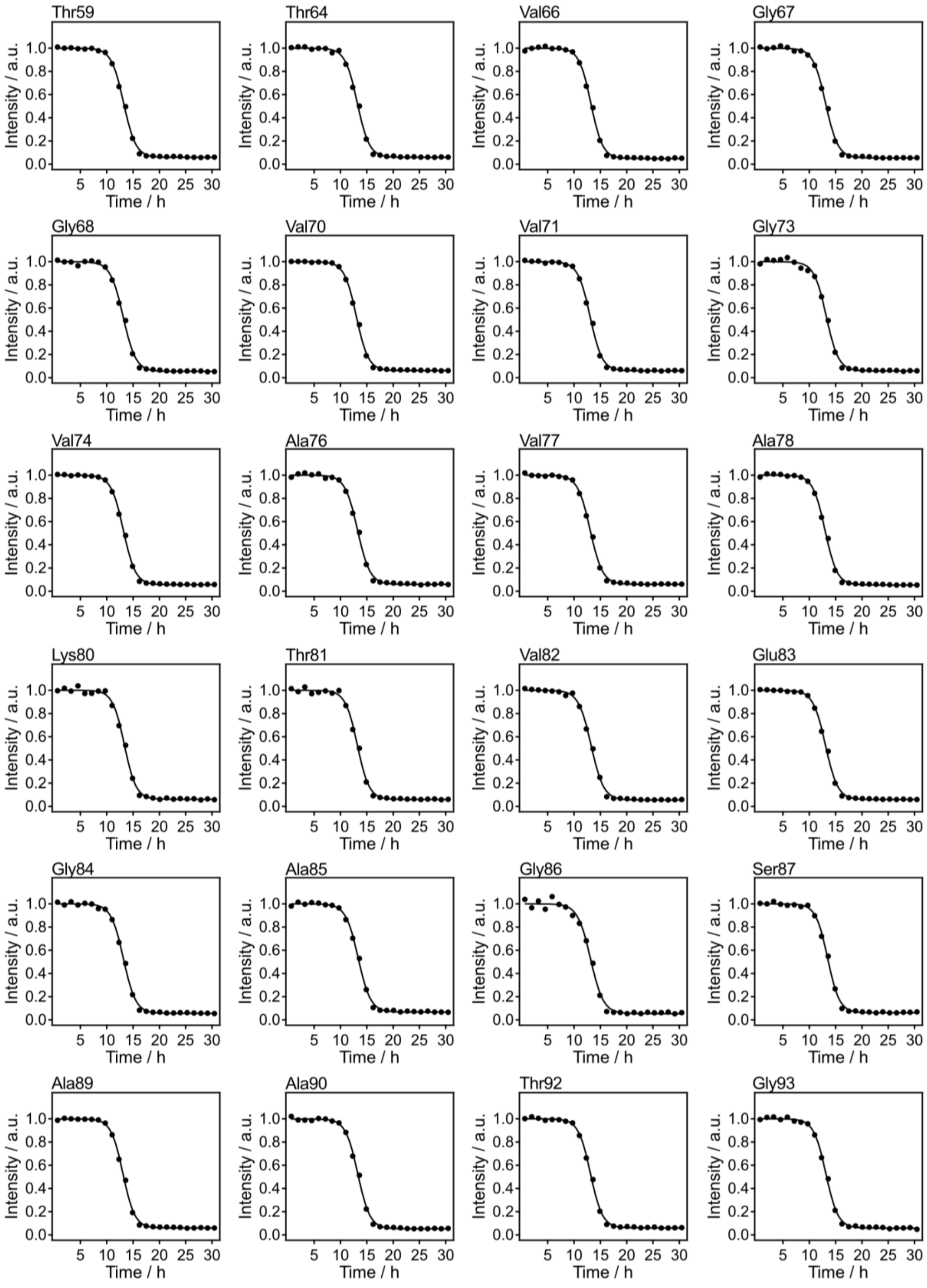


**Extended Data Fig. 4 | Rheo-NMR measurements of αSyn amyloid fibril formation under dilute conditions.** Time courses of $^{1}H$-$^{15}N$ HSQC peak intensities for assigned residues from Thr59 to Gly93 during 20-Hz rotation at 30 °C without PEG. Intensities were normalized to the values before rotation. Solid lines indicate the best-fit curves obtained using the equation shown in Methods. Error bars represent fitting errors.

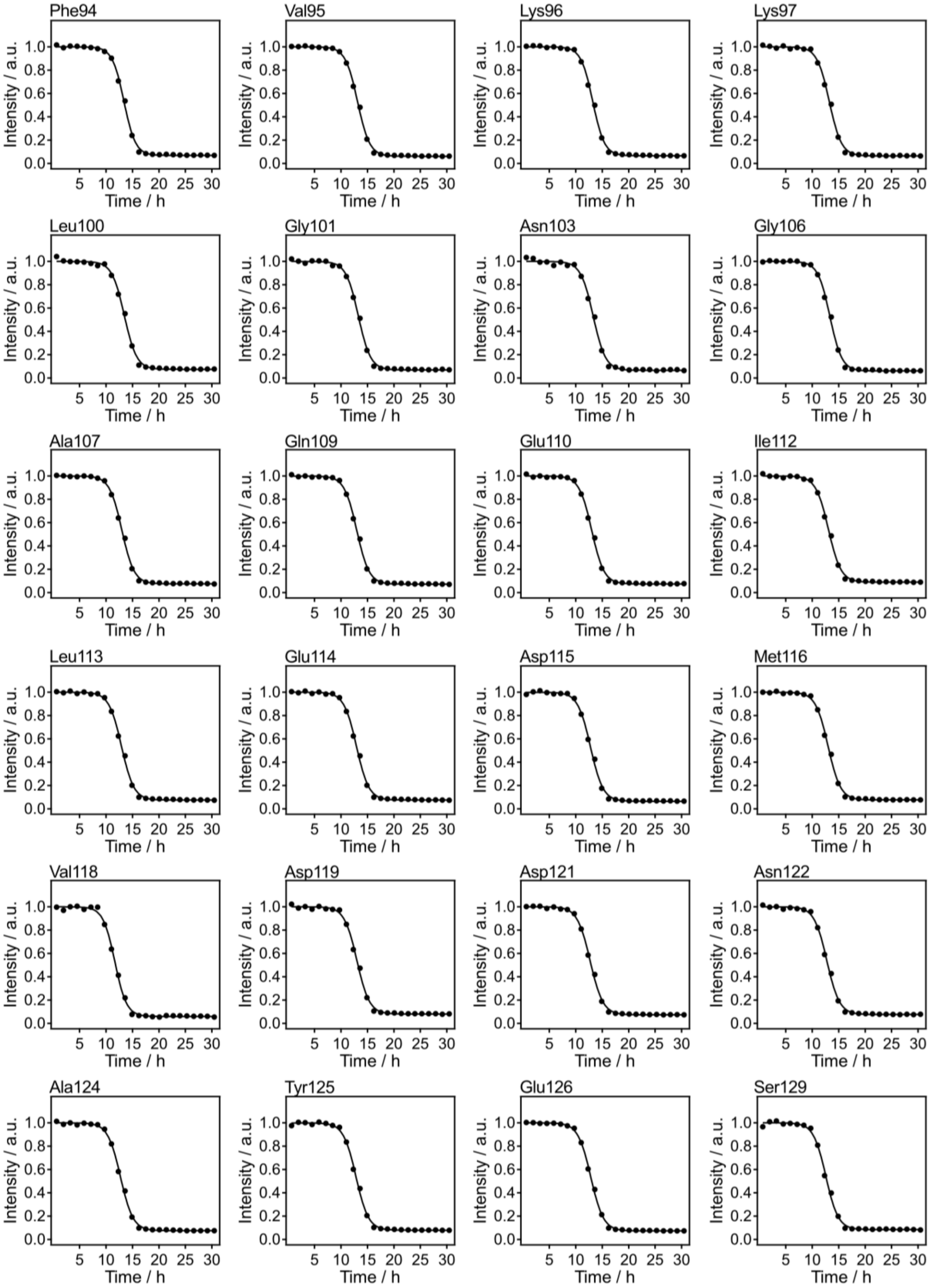


**Extended Data Fig. 5 | Rheo-NMR measurements of αSyn amyloid fibril formation under dilute conditions.** Time courses of $^{1}H$-$^{15}N$ HSQC peak intensities for assigned residues from Phe94 to Ser129 during 20-Hz rotation at 30 °C without PEG. Intensities were normalized to the values before rotation. Solid lines indicate the best-fit curves obtained using the equation shown in Methods. Error bars represent fitting errors.

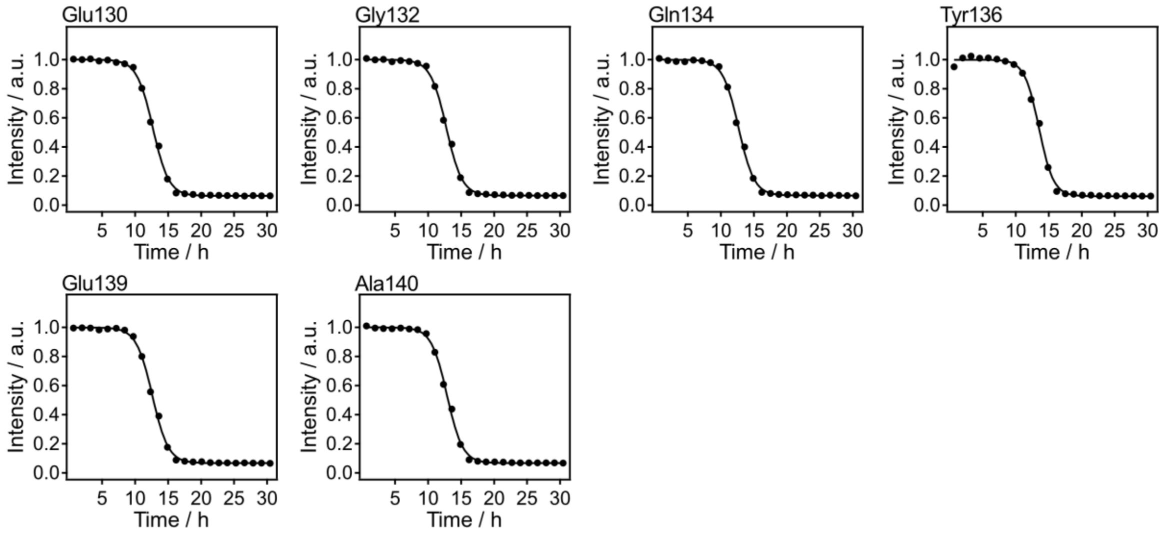


**Extended Data Fig. 6 | Rheo-NMR measurements of αSyn amyloid fibril formation under dilute conditions.** Time courses of $^{1}$H-$^{15}$N HSQC peak intensities for assigned residues from Glu130 to Ala140 during 20-Hz rotation at 30 °C without PEG. Intensities were normalized to the values before rotation. Solid lines indicate the best-fit curves obtained using the equation shown in Methods. Error bars represent fitting errors.

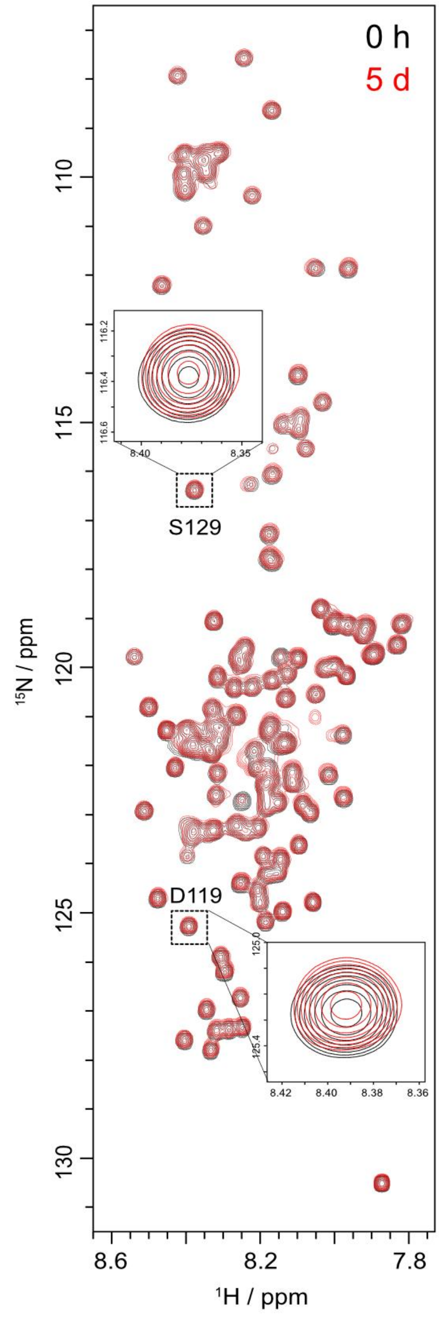


**Extended Data Fig. 7 | NMR spectra in the presence of PEG under quiescent conditions.** $^{1}H$-$^{15}N$ HSQC spectra of αSyn in the presence of 10% (w/v) PEG 8000 before (black) and after 5 days (red) of quiescent incubation at 25 °C. Expanded views of the signals from Asp119 and Ser129 are shown.

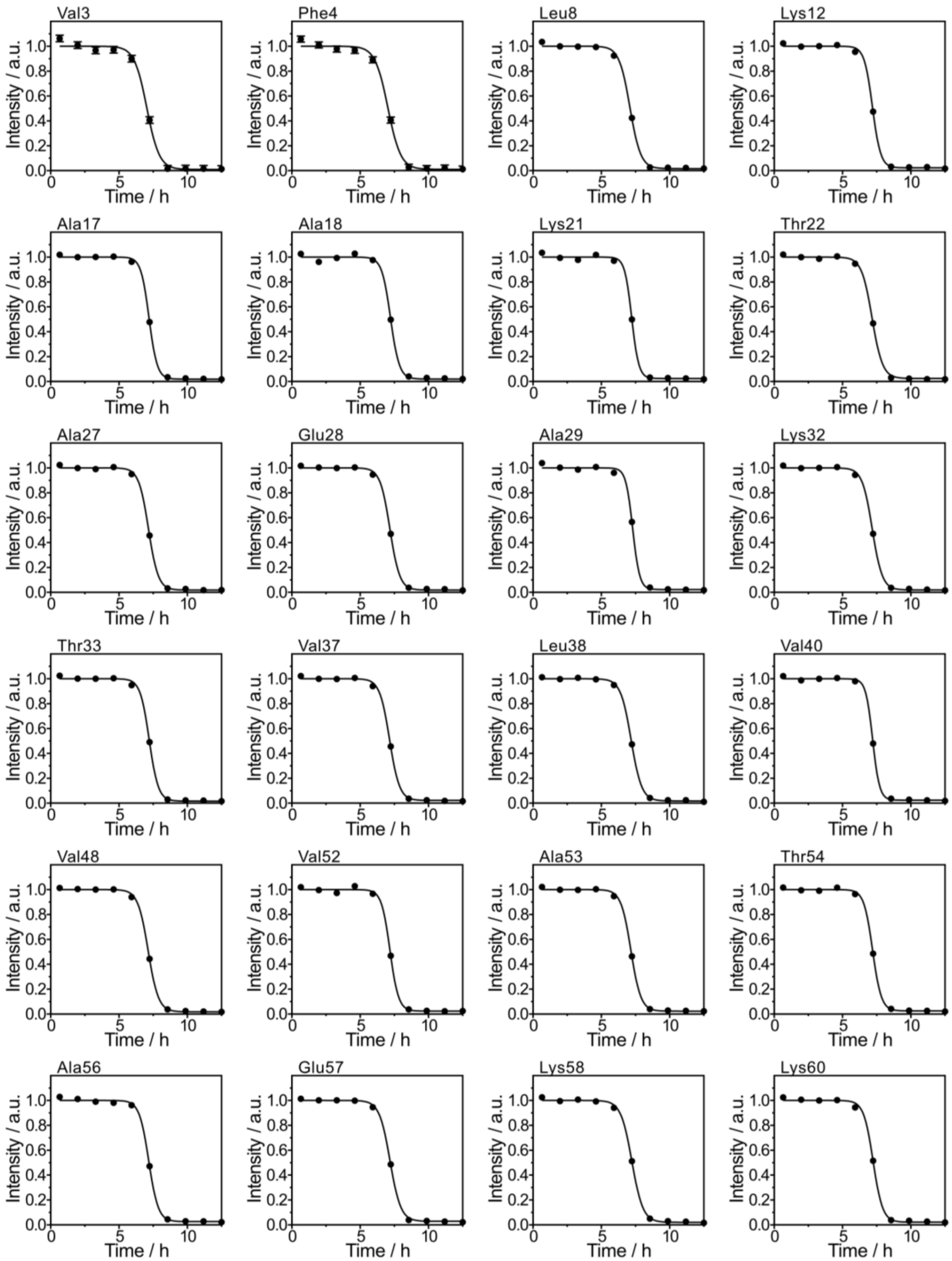


**Extended Data Fig. 8 | Rheo-NMR measurements of αSyn amyloid fibril formation under PEG-induced LLPS conditions.** Time courses of $^{1}H$-$^{15}N$ HSQC peak intensities for assigned residues from Val3 to Lys60 during 20-Hz rotation at 25 °C with 10% (w/v) PEG 8000. Intensities were normalized to the values before rotation. Solid lines indicate the best-fit curves obtained using the equation shown in Methods. Error bars represent fitting errors.

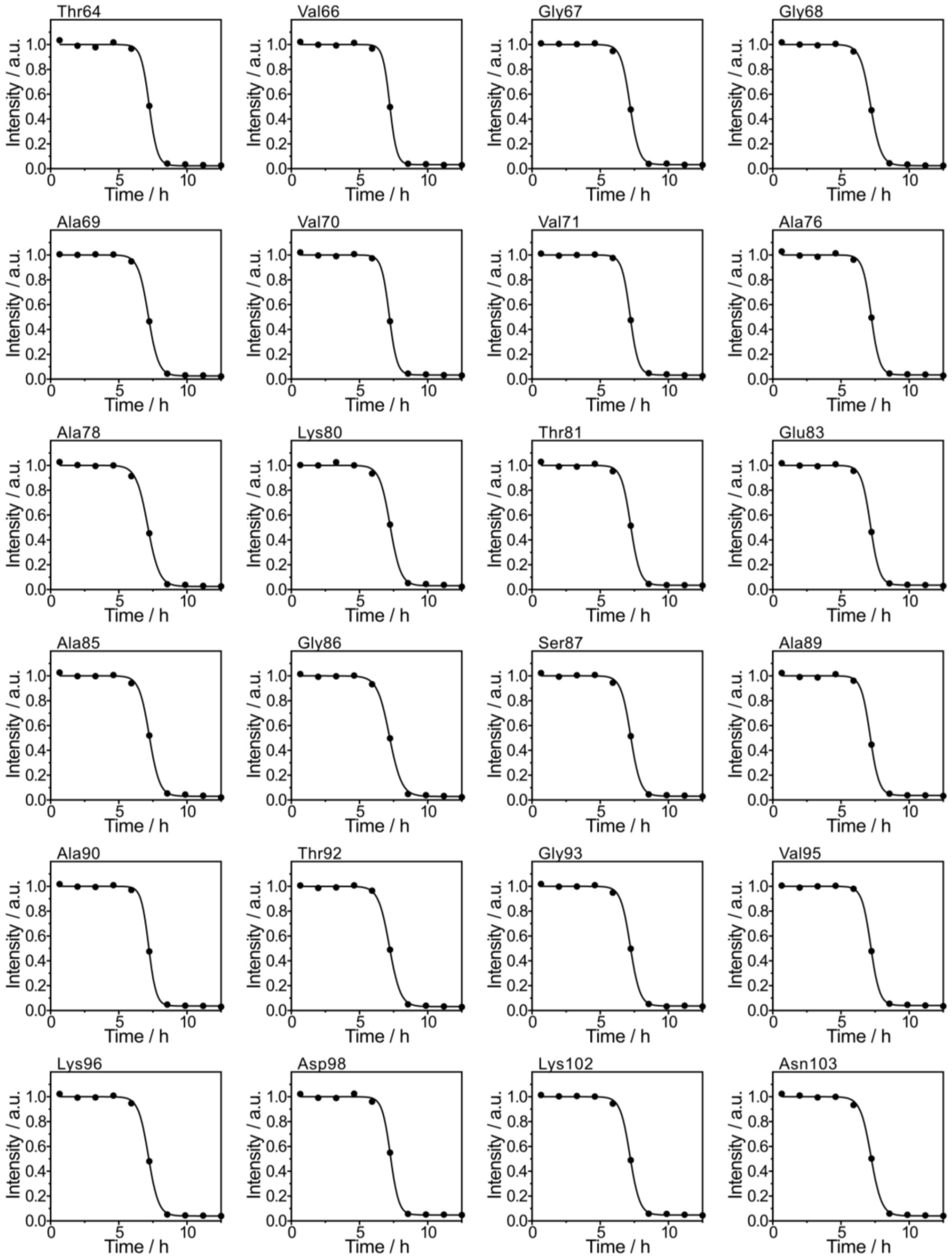


**Extended Data Fig. 9 | Rheo-NMR measurements of αSyn amyloid fibril formation under PEG-induced LLPS conditions.** Time courses of $^{1}H$-$^{15}N$ HSQC peak intensities for assigned residues from Thr64 to Asn103 during 20-Hz rotation at 25 °C with 10% (w/v) PEG 8000. Intensities were normalized to the values before rotation. Solid lines indicate the best-fit curves obtained using the equation shown in Methods. Error bars represent fitting errors.

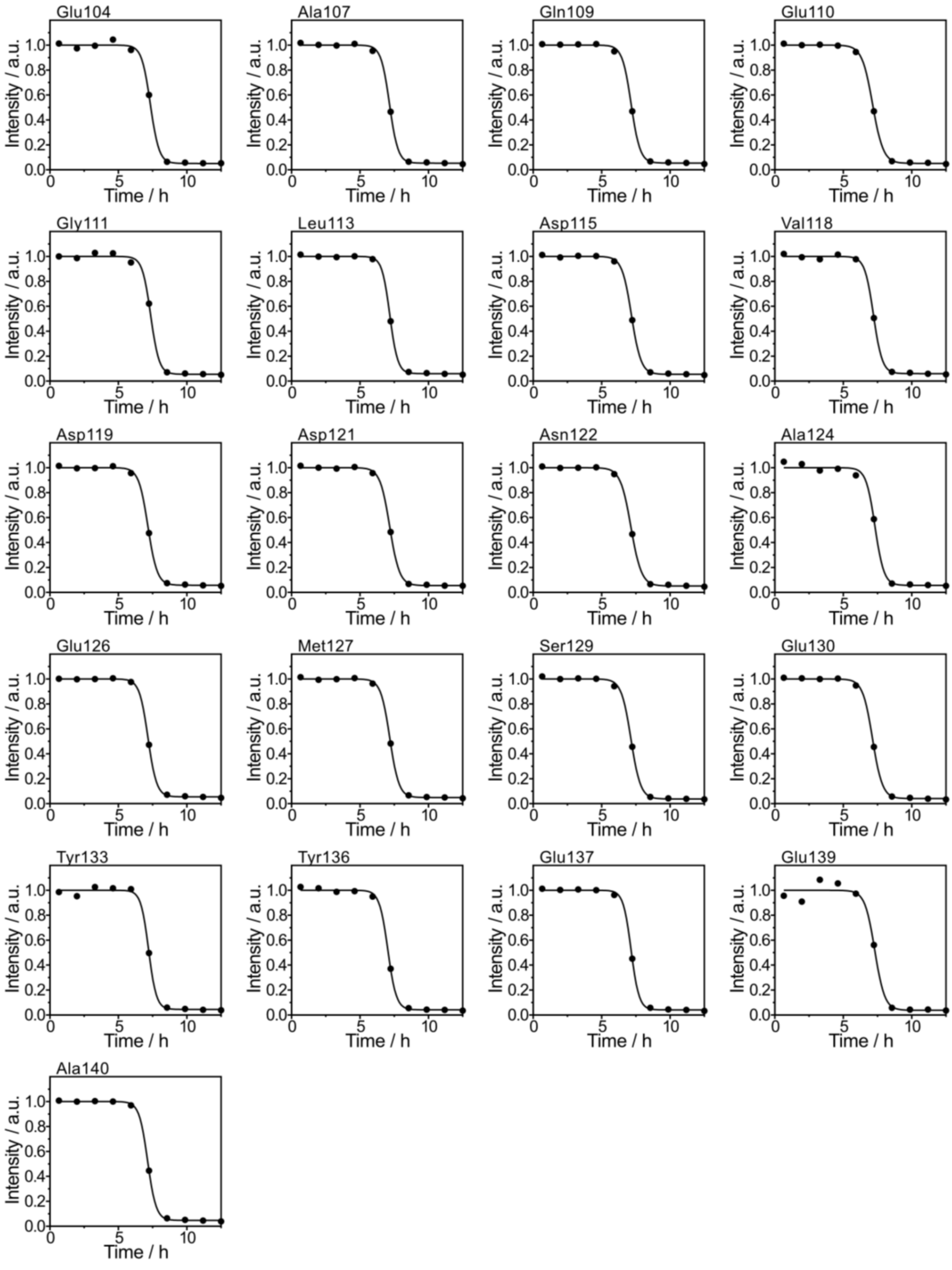


**Extended Data Fig. 10 | Rheo-NMR measurements of αSyn amyloid fibril formation under PEG-induced LLPS conditions.** Time courses of $^{1}H$-$^{15}N$ HSQC peak intensities for assigned residues from Glu104 to Ala140 during 20-Hz rotation at 25 °C with 10% (w/v) PEG 8000. Intensities were normalized to the values before rotation. Solid lines indicate the best-fit curves obtained using the equation shown in Methods. Error bars represent fitting errors.
